# *C. elegans* CED-1 acts in neurons to modulate ciliary protein abundance and extracellular vesicle shedding

**DOI:** 10.1101/2025.09.15.676301

**Authors:** Tao Ke, Jessica E. Tanis

## Abstract

The *C. elegans* receptor CED-1, along with orthologs mammalian MEGF10 and *Drosophila* Draper, plays a well-established and conserved role in phagocytosis by acting in engulfing cells.^1–7^ Interestingly, CED-1 family members are also expressed in neurons, but their functions in these non-engulfing cells remain unclear.^6,8,9^ Our study shows that CED-1 localizes to the dendrites and primary cilia of male tail RnB neurons, which mediate sensory perceptions during mating^10^ and generate extracellular vesicles (EVs) that transfer bioactive macromolecules for both intercellular and animal-to-animal communication.^11^ Loss of *ced-1* leads to a reduction in the shedding of EVs that contain the transient receptor potential (TRP) channel PKD-2 from the cilium distal tip, and this is rescued by the expression of CED-1 in the neurons. CED-1 is required to increase both the abundance of PKD-2 in the cilium and PKD-2 EV shedding in response to the physiological stimulus of mating partners. Assessment of *ced-1* mutant male mating indicates that neuronal CED-1 is also important for turning behavior, which helps the male tail to maintain contact with a mate. Collectively, these results reveal a new role for CED-1 in neurons as a regulator of EV biogenesis in response to environmental cues, optimizing the shedding of bioactive EVs for effective communication.

## RESULTS

The phagocytic receptor *C. elegans* CED-1 acts in engulfing cells to enable recognition and removal of apoptotic cell corpses during development,^6^ axon fragments following neuronal injury,^7^ large vesicles termed exophers generated by neuronal stress,^12^ and vesicles shed from neuronal processes that contain hyperactive kinesin-2.^13^ Despite the well-established roles for CED-1 and its orthologs, mammalian MEGF10 and Drosophila Draper, in phagocytosis,^1–7,12–14^ the function of these proteins in non-engulfing cells remains poorly understood. In this study, we define a new neuronal role for CED-1 in the regulation of extracellular vesicle biogenesis, ciliary protein localization, and male mating behavior.

### CED-1 is expressed in male tail RnB neurons

CED-1 was reported to be expressed in *C. elegans* neurons, though these data were never shown.^6^ To define where CED-1 localizes to in neurons, we inserted a C-terminal mNeonGreen tag at the *ced-1* endogenous locus (CED-1::mNG). Fusion of GFP to CED-1 at this position results in a functional protein that rescues *ced-1* mutant engulfment defects.^6^ We observed CED-1::mNG expression in the male tail sensory rays (Figure 1A, 1B, and Figure S1) required for mating behavior,^10^ in addition to hypodermal, muscle, and gonadal sheath cells as previously described.^6^ Analysis of CED-1::mNG in animals expressing mCherry from the *col-10* promoter,^15^ showed that CED-1 localized not only to hypodermal cells in the male tail, but also other cells in the rays (Figure S1). To define these cells, we used the *klp-6* promoter to drive mCherry expression in male tail ciliated sensory neurons.^16^ Colocalization analysis revealed that CED-1::mNG is expressed in the dendrites and ciliary region of B-type ray neurons (RnB) (Figure 1A and 1C).

**Figure 1.**
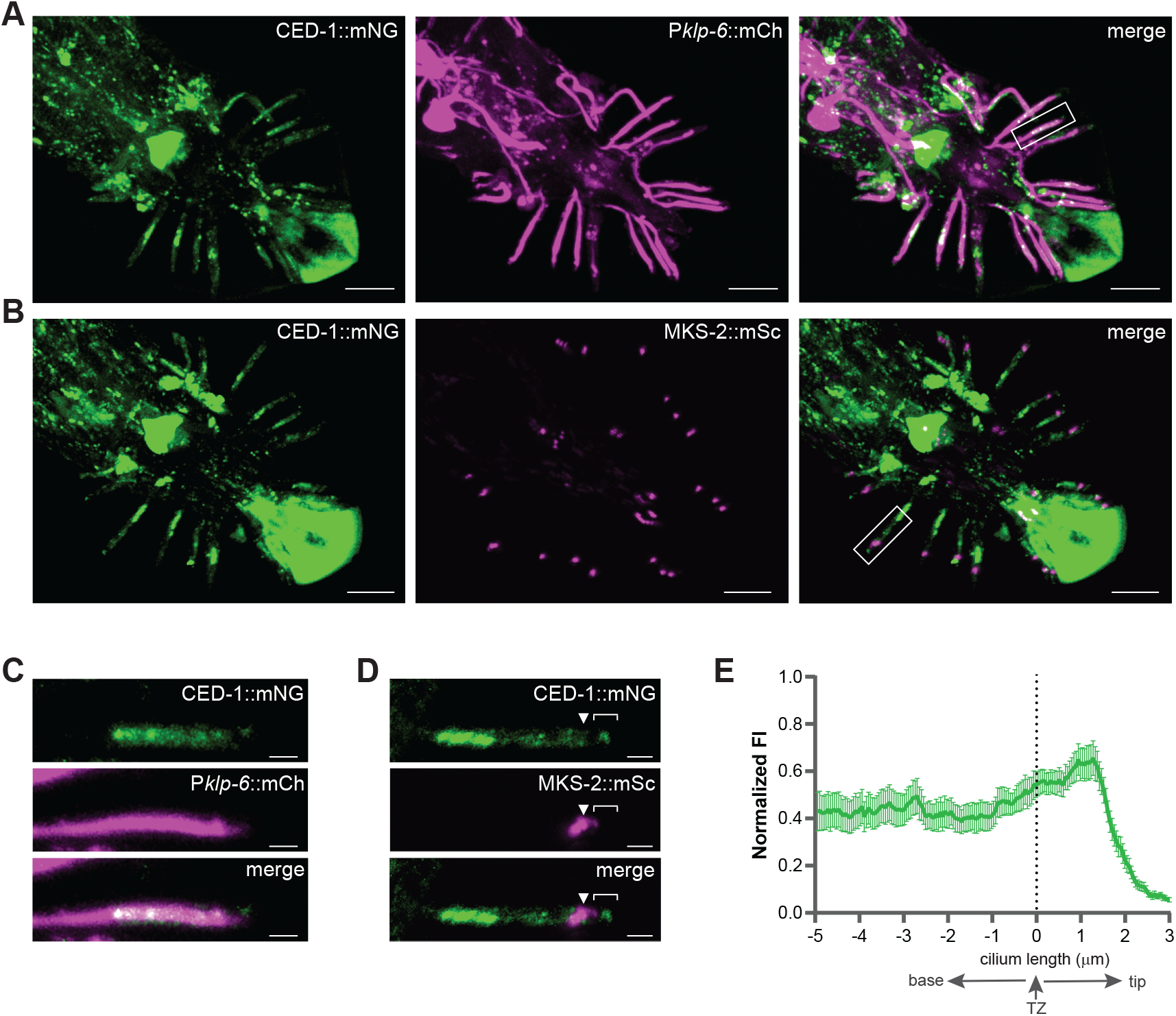
CED-1 localizes to the RnB neurons in the male tail. (A) Coexpression of a CED-1::mNG endogenous reporter (left) with a P*klp-6*::mCh transgene (middle) shows that CED-1 is expressed in RnB and HOB EVNs as well as other cells in the male tail (merge, right). (B) Coexpression of CED-1::mNG (left) with MKS-2::mSc to label the TZ (middle). In A and B, scale bars, 10 µm. (C) CED-1::mNG is present in the dendrite and ciliary region in a representative R4B neuron (box, right panel A); scale, 2 µm. (D) In relation to the MKS-2::mSc-labeled TZ, CED-1 localizes to the cilium proper, base and dendrite (box, right panel B). Cilium proper ([), transition zone (▾), ciliary base ({), and scale, 2 μm. (E) Normalized fluorescence intensity of CED-1::mNG along dorsal R1B and R5B cilia. Data are represented as mean ± SEM (n = 16); center of the TZ indicated with dotted line.

We next sought to more clearly define the localization of this protein in the RnB neurons. The dendritic endings of the RnBs terminate in primary cilia, which are enriched in receptors that receive signals from the environment to mediate sensory perception during mating.^17^ The transition zone (TZ) acts as a molecular gate to permit selective transport of specific proteins from the ciliary base into the cilium proper.^18^ To determine where CED-1 localizes, we co-expressed CED-1::mNG with mScarlet-tagged MKS-2 (MKS-2::mSc), which labels the TZ (Figure 1B and 1D).^19,20^ Lengthwise profiling of fluorescent signal in the RnB neurons showed CED-1 in the distal dendrite, ciliary base, and cilium proper (Figure 1E).

### Loss of *ced-1* reduces shedding of extracellular vesicles from the cilium distal tip

*C. elegans* ciliated neurons transmit signals through the shedding of extracellular vesicles (EVs) termed ectosomes, which mediate intercellular and inter-organismal transport of bioactive macromolecules to control behavior.^11,21–25^ Distinct EV subpopulations shed from the RnB neurons are released through a pore in the cuticle into the environment (Figure 2A). EVs containing the CLHM-1 ion channel^26^ bud from the periciliary membrane compartment (PCMC) at the ciliary base, while EVs containing the TRP ion channel PKD-2 are predominantly shed from the cilium distal tip.^24,25^ To determine if CED-1 impacts EV biogenesis, we created a strain containing the *ced-1(e1735)* early stop mutation, tdTomato-tagged CLHM-1 (CLHM-1::tdT) and GFP-tagged PKD-2 (PKD-2::GFP) single copy transgenes,^25^ and *him-5(e1490)*, which increases the frequency of male progeny due to X chromosome nondisjunction.^27^ We imaged EVs released into the environment from male tails using total internal reflection fluorescence (TIRF) microscopy (Figure 2B, Figure S2). Loss of *ced-1* significantly reduced PKD-2::GFP EV release, but had no effect on the shedding of CLHM-1-containing EVs from the ciliary base or the colocalization of the ion channel cargos in EVs (Figure 2C, 2D, and 2E). This suggests that CED-1 influences the shedding of EVs from cilium tip where ectosomes are most abundantly generated.

**Figure 2.**
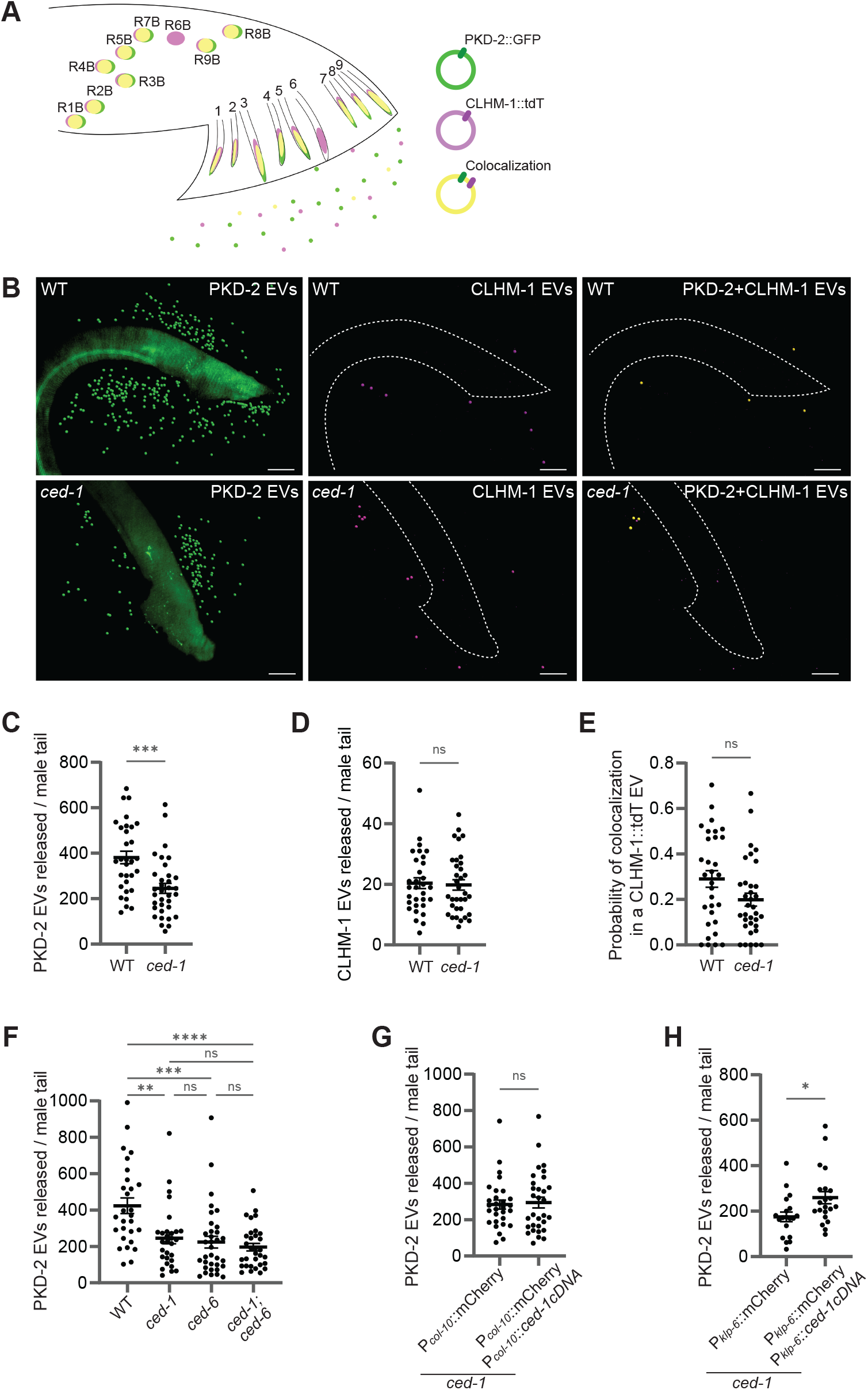
Neuronal CED-1 regulates EV shedding from the cilium distal tip. (A) Schematic of EVs released from *C. elegans* RnB cilia into the environment. (B) Representative images of PKD-2::GFP (*henSi21*) and CLHM-1::tdT (*henSi3*) EVs released from wild type (top) and *ced-1(e1735)* mutant (bottom) animals; scale, 15 μm. EVs are marked with spots (Imaris software) for visualization; original images are in Figure S2. (C) PKD-2::GFP EV release is significantly reduced in *ced-1* mutants; n ≥ 31. (D) CLHM-1::tdT EV release and (E) probability of PKD-2::GFP colocalization in a CLHM-1::tdT EV is unaffected by loss of *ced-1*; n ≥ 31. (F) *ced-6(hen38)* mutant males exhibit reduced PKD-2::GFP EV release. No further decrease is observed in *ced-1; ced-6* double mutants; n ≥ 28. (G) Expression of *ced-1* in the hypodermis (*col-10* promoter) has no effect on PKD-2::GFP EV shedding in *ced-1* mutants; n ≥ 19. (H) *ced-1* expression in RnB and HOB neurons (*klp-6* promoter) is sufficient to increase PKD-2::GFP EV release in the *ced-1* mutant; n ≥ 19. In C-H, data are represented as mean ± SEM. For (F), a one-way ANOVA and Turkey’s multiple comparisons test was performed; a Mann-Whitney test was used for all other analyses; *<p<0.05, **p<0.01, ***p<0.001, ****p<0.0001.

The adaptor protein CED-6 interacts with the C-terminus of CED-1 to mediate removal of apoptotic cells and clearance of axonal debris such that *ced-1; ced-6* double mutants exhibit engulfment defects indistinguishable from either single mutant.^6,7^ To determine if CED-6 functions with CED-1 to regulate EV biogenesis, we compared the number of PKD-2::GFP EVs shed from the *ced-6(hen38)* deletion mutant and *ced-1; ced-6* double mutant to *ced-1* mutant and control animals (Figure 2F). Loss of *ced-6* resulted in release of significantly fewer PKD-2 EVs compared to the control. However, there was no further decrease in the number of PKD-2 EVs shed by the *ced-1; ced-6* animals compared to the single mutants, suggesting that CED-1 and CED-6 function together to modulate PKD-2 EV biogenesis.

### CED-1 functions in RnB neurons to regulate EV release in response to mates

Cell-specific rescue experiments have demonstrated CED-1 function in engulfing cells. Hypodermal expression of *ced-1* is sufficient for engulfment of somatic cell corpses, exophers, and axon fragments,^6,12,28^ gonadal sheath expression leads to engulfment of germ cell corpses^6^, and muscle expression promotes axon regeneration and removal of axonal debris.^7^ Further, CED-1 function in the hypodermis is required for efficient production of exophers by mechanosensory neurons.^12^ To determine where CED-1 acts to regulate EV biogenesis, we expressed the *ced-1* cDNA from either the *col-10* hypodermal promoter^15^ or *klp-6* EV-releasing neuron (EVN) promoter^16^ in the *ced-1* mutant and quantified PKD-2::GFP EV release. Compared to the *ced-1* mutant control, expression of *ced-1* in the hypodermis did not affect the shedding of PKD-2 EVs (Figure 2G). Surprisingly, expression of *ced-1* in the EVNs increased the number of PKD-2 EVs released from *ced-1* mutant animals (Figure 2H). Our results indicate that CED-1 function in the RnB neurons is sufficient to regulate shedding of cilium-tip derived EVs.

Males transfer PKD-2-containing EVs to the hermaphrodite vulva during copulation^23^ and the presence of mating partners impacts EV cargo composition and release into the environment,^24^ suggesting that EVs are important for inter-organismal communication. To determine if loss of *ced-1* impacts EV shedding in response to mates, we quantitated EV release from virgin and mated males. Control males raised with hermaphrodites released significantly more PKD-2::GFP EVs than virgin males, however, *ced-1* mutants failed to increase EV shedding in the presence of mating partners (Figure 3A). This suggests that CED-1 is required to modulate EV biogenesis in response to hermaphrodites.

**Figure 3.**
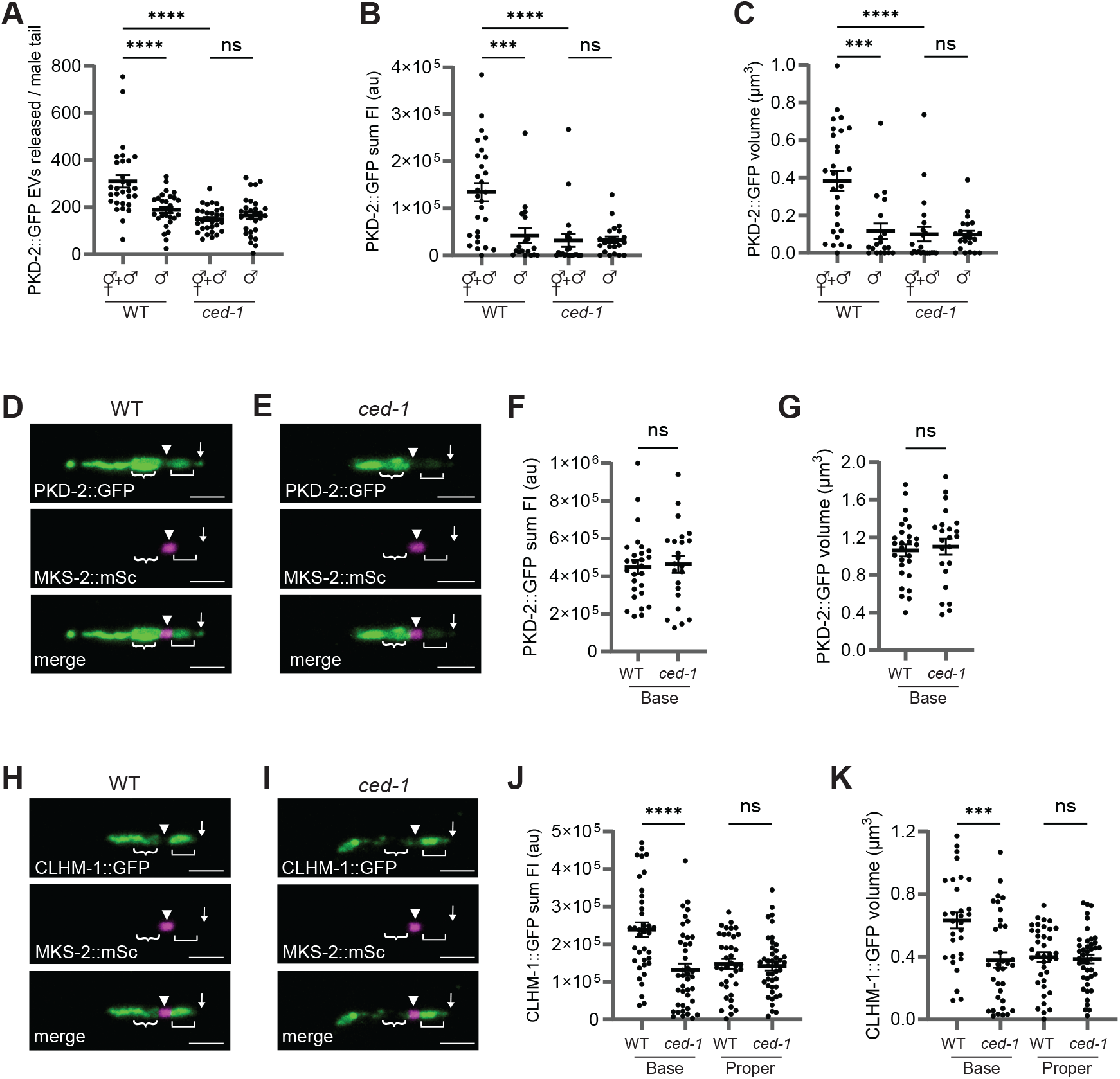
CED-1 and the presence of mating partners affect the abundance of ciliary cargos in EVs. (A) Compared to virgin males, release of PKD-2 EVs from males raised with mating partners is increased in WT, but not *ced-1* mutant animals; n ≥ 29. (B) Virgin males exhibit a decrease in PKD-2::GFP fluorescence intensity and (C) volume in the cilium proper compared to males grown with hermaphrodites. Loss of *ced-1* eliminates the impact of mating partners on PKD-2::GFP enrichment in the cilium; n ≥ 18. (D) Representative images of PKD-2::GFP (top) and MKS-2::mSc (middle) in WT and (E) *ced-1* mutant RnBs. (F) Loss of *ced-1* does not alter PKD-2::GFP fluorescence intensity or (G) volume in the ciliary base n ≥ 23. (H) Representative images of CLHM-1::GFP (top) and MKS-2::mSc (middle) in WT and (I) and *ced-1* mutant RnBs. (J) Loss of *ced-1* causes a decrease in CLHM-1::GFP fluorescence intensity and (K) volume in the ciliary base, but not the cilium proper n ≥ 30. All images are labeled as cilium distal tip (↓), cilium proper ([), transition zone (▼), ciliary base ({), and scale, 2 μm. Data are mean +/−SEM. For A-C, a one-way ANOVA with Turkey’s multiple comparisons test was used; a Mann-Whitney test was used for other analyses (***p<0.001, ****p<0.0001).

### CED-1 affects ciliary abundance of PKD-2

Shedding of PKD-2::GFP labeled EVs from the cilium tip is a dynamic process that is dependent on the presence of PKD-2 in the cilium^24,25^ and replenishment of PKD-2 at the distal tip.^29^ Quantification of PKD-2::GFP in the cilium revealed that PKD-2 fluorescence intensity and volume was significantly lower in virgin males compared to males raised with hermaphrodites (Figure 3B and 3C), suggesting that the presence of mating partners impacts cilium enrichment of PKD-2. We next sought to determine how loss of *ced-1* affects the ciliary pool of PKD-2. Compared to wild type, *ced-1* mutants raised with mating partners exhibited significantly less PKD-2::GFP in the cilium (Figure 3B, 3C, 3D, and 3E). Further, there was no difference in PKD-2 fluorescence intensity or volume in *ced-1* mutants raised with or without hermaphrodites (Figure 3B and 3C). PKD-2::GFP abundance in the ciliary base was unaffected in *ced-1* mutants (Figures 3F and 3G). Together, these results suggest that loss of *ced-1* prevents an increase in ciliary PKD-2 in response to mating partners, which could underlie the reduced PKD-2 EV shedding observed in the *ced-1* mutant.

Next, we investigated whether CED-1 was required for proper ciliary abundance of a different EV cargo, CLHM-1 (Figures 3H and 3I). Despite having no effect on release of CLHM-1 EVs into the environment, loss of *ced-1* significantly reduced CLHM-1 in the ciliary base, but not the cilium proper (Figures 3J and 3K). This is consistent with a role for CED-1 in the enrichment of ciliary proteins in different compartments.

### CED-1 regulates male turning behaviors of mating males

When a male *C. elegans* encounters a mating partner, he immediately responds by pausing forward movement and pressing his tail against the hermaphrodite in order to search for the vulva.^10^ Upon reaching the hermaphrodite head or tail, he must turn to scan on the other side of the body (Figure 4A). Once he reaches the vulva, he stops and coordinates his movement with that of the hermaphrodite in order to successfully mate.^10^ The RnB neurons and PKD-2 control multiple aspects of mating.^30–32^ To determine if the *ced-1* mutant has defects in this complex behavior, we quantified response to hermaphrodites, ventral touch time, number of successful versus failed turns, and vulva contact duration (Figures 4B, 4C, 4D and Table S1). While *ced-1* mutants did not exhibit defects in response or location of vulva behaviors (Table S1), the absence of *ced-1* increased the number of failed turns, thus reducing the percent of successfully completed turns compared to the control (Figures 4C and 4D). As activation of the RnB neurons is sufficient to induce ventral curling of the male tail,^32^ which could impact turn completion rate, we sought to determine if CED-1 function in the EVNs is sufficient to affect this behavior. Expression of the *ced-1* cDNA from the *klp-6* promoter significantly increased the percent of successful turns in the *ced-1* mutants (Figure 4E, 4F, and 4G). These results demonstrate that CED-1 function in the RnB neurons is sufficient to modulate male turning behavior during mating.

**Figure 4.**
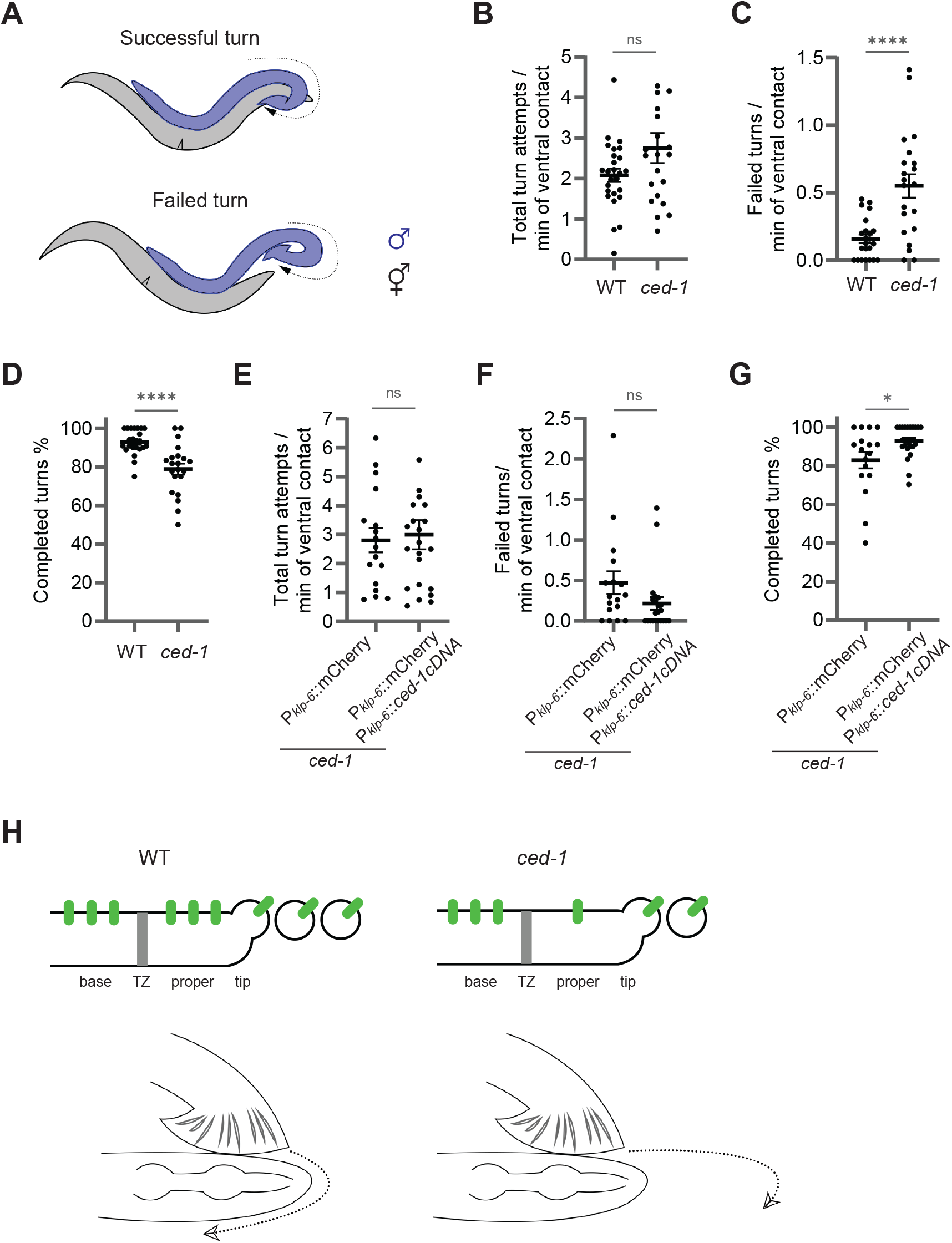
CED-1 regulates male turning behavior during mating. (A) Schematic of male turning, while interacting with a hermaphrodite mate. (B) The total number of turn attempts per minute of hermaphrodite contact time was not different in *ced-1(e1735)* males compared to the control. (C) Loss of *ced-1* increased the number of failed turns and (D) reduced the percent of successfully completed turns. In B-D, n ≥ 21. (E) Expression of the *ced-1* cDNA in the EVNs (*klp-6 promoter*) of the *ced-1* mutant did not affect the total number of turn events, but (F) caused a downward trend in the number of failed turns (p=0.103) and (G) significantly increased the percent of successfully completed turns n ≥ 17. Data are represented as mean ± SEM; Mann-Whitney test used for analysis, *p<0.05, ****p<0.0001. (H) Model; CED-1 functions in the RnB neurons to regulate ciliary abundance of PKD-2, shedding of EVs from the cilium distal tip, and male turning during contact with mating partners.

## DISCUSSION

EVs play key roles in physiological processes as well as the propagation of pathophysiological conditions by enabling bioactive macromolecules that cannot cross the plasma membrane to be shared between cells.^33–36^ We identified CED-1 as a new regulator of EV biogenesis by using *C. elegans* with fluorescent protein-tagged EV cargoes to visualize ectosomes released from neuronal primary cilia. Loss of *ced-1* reduced the shedding of PKD-2-containing EVs from the cilium distal tip, and this was positively correlated with the ciliary abundance of this EV cargo. While previously recognized for its function in engulfing cells,^6,7,12,13^ our findings demonstrate that CED-1 acts in neurons to modulate EV shedding in response to the presence of mating partners. As CED-1 orthologs MEGF10 and Draper are expressed in neurons,^8,9^ this raises the possibility that these conserved family members could also mediate neuronal EV release in response to physiological cues.

How might CED-1 regulate PKD-2 ciliary abundance and EV shedding? Early during the engulfment of apoptotic cell corpses, the activation of CED-1 leads to actin reorganization within phagosomes of engulfing cells.^37^ Although cilia are microtubule-based organelles, the actin cytoskeleton plays a crucial role in many ciliary processes, including ciliogenesis, intraflagellar transport (IFT), cilium tip ectocytosis, and cilium disassembly.^38–45^ Actin polymerization directly mediates the release of ectosomes from the cilium distal tip.^44^ It is possible that the loss of *ced-1* affects the dynamic remodeling of actin filaments necessary for the efficient release of ciliary EVs, leading to a decrease in shedding of PKD-2 EVs into the environment. However, it is important to note that a direct impact of CED-1 on ectocytosis would not account for the reduced abundance of PKD-2 in the cilium observed in the *ced-1* mutants.

For optimal detection of signals and EV output, the levels of proteins in different ciliary compartments are controlled by IFT.^21,24,25^ In mammalian primary cilia, the lysine demethylase KDM3A regulates IFT in the cilium and periciliary compartment by modulating actin dynamics.^38^ The actin cytoskeleton also controls the entry of IFT trains into flagella in *Chlamydomonas reinhardtii*.^40^ We propose that, in the presence of a signal from mating partners, activation of CED-1 could alter actin dynamics to increase IFT of PKD-2 into the cilium. This would enable males to accumulate sufficient ciliary PKD-2 to package into EVs that are deposited on vulva of hermaphrodites during mating. ^23^

CED-1 and CED-6 act non-cell autonomously to clear severed axons and initiate axon regeneration.^7,28^ CED-1 homologs MEGF10 and Draper also impact neuron morphology and function by acting in engulfing cells.^1–5,14^ While MEGF10 is expressed in cholinergic and glutamatergic neurons in the brain,^8^ the only previously known neuronal function for MEGF10 is in defining the spatial arrangement of starburst amacrine cells (SACs) and horizontal cells in the retina.^9,46^ These interneuron subtypes form mosaic arrays such that it is unlikely for two neurons of the same subtype to be adjacent to each other, resulting in an even distribution of these cells across the retina.^47^ The establishment of SAC mosaics is dependent on a MEGF10-mediated repulsive signal between the dendritic arbors and the somas of nearby homotypic neurons, as SACs with conditional knockout of MEGF10 are unable to repel neighboring SACs that express functional MEGF10.^46^ The exact mechanism underlying this repulsion has yet to be elucidated. SACs produce EVs, and loss of the *Pten* phosphatase alters the proteome of retinal EVs, increases internalization of MEGF10, and disrupts SAC patterning.^48^ Our findings indicate that CED-1 can regulate both the abundance of EV cargos and the number of EVs released. This raises the intriguing possibility that MEGF10 may regulate the biogenesis of EVs that carry repulsive cues necessary for forming neuronal mosaics.

## SUPPLEMENTAL INFORMATION

Supplemental information includes two figures and one table and can be found with this article online.

## ACKNOWLEDGMENTS

We thank the *Caenorhabditis* Genetics Center, which is supported by National Institutes of Health (NIH) ORIP P40 OD010440, for strains. Microscopy access was supported by NIH-NIGMS P20 GM103446, NIGMS P20 GM139760, and the State of Delaware; equipment was acquired with NIGMS S10 OD030321. A UD BioImaging Core Center Access Voucher (to T.K.) was used to cover some imaging expenses. This work was supported by NIH-NIGMS R01 GM135433 (to J.E.T.).

## AUTHOR CONTRIBUTIONS

Research Design, T.K. and J.E.T.; Methodology, T.K. and J.E.T.; Data Collection, T.K.; Data Analysis, T.K.; Writing – original draft, T.K. and J.E.T.; Writing – review and editing, T.K. and J.E.T.; Visualization, T.K.; Funding Acquisition, T.K. and J.E.T.

## DECLARATION OF INTERESTS

The authors declare no competing interests.

## FIGURE LEGENDS

**Supplemental Figure S1**. CED-1 is expressed in the hypodermis in the male tail. (A) Coexpression of the CED-1::mNG endogenous reporter (left) with a P*col-10*::mCherry (middle). Bottom, merge; scale bars, 10 µm.

**Supplemental Figure S2**. CED-1 impacts EV shedding from male tail EVNs. Representative images of PKD-2::GFP (*henSi21*) and CLHM-1::tdT (*henSi3*) labeled EVs released from WT (left) and *ced-1(e1735)* mutant males. Images correspond to those in Fig. 2B, scale bars, 10 µm.

**Table S1**. Mating behaviors of WT and *ced-1(e1735)* males

## STAR METHODS

## RESOURCE AVAILABILITY

### Lead contact

Further information and reagent requests should be directed to and will be fulfilled by the lead contact, Jessica Tanis

### Materials availability

*C. elegans* strains generated in the study will be shared upon request. The University of Delaware Material Transfer Agreement Request Webform will be completed when research materials are transferred to an outside party.

### Data and code availability

All original data have been deposited at Mendeley Data and are publicly available as of the data of publication

This study does not report original code

Any additional information required to reanalyze the data reported in this paper is available from the lead contact upon request.

## EXPERIMENTAL MODEL AND SUBJECT DETAILS

### *C. elegans* strains and maintenance

All strains were maintained at 20°C on Nematode Growth Media (NGM) plates with OP50 *E. coli. ced-1(e1735)* I, *ced-6(hen38)* III, and *him-5(e1490)* V are mutant alleles. *ced-1(syb9348* [*ced-1::mNG*]) I and *mks-2(syb7299* [*mks-2::mSc*]) II are endogenous insertion alleles. *henSi3* [*clhm-1 promoter::clhm-1::tdT*] III, *drSi33* [*clhm-1 promoter::clhm-1::GFP*] IV, and *henSi21* [*pkd-2 promoter::pkd-2::GFP*] V are single copy transgenes. All strains used for analysis of EV release, ciliary abundance, and male mating behavior contain *him-5(e1490)*; the “WT” control has *him-5(e1490)*, but no other mutations. Duplex genotyping was used to detect *ced-6(hen38)*; SuperSelective genotyping was used to detect *ced-1 (e1735)*.^49^ See Key Resources Table for all strains and genotyping oligonucleotides used in this study. Some strains are available from the *Caenorhabditis* Genetics Center (CGC); all other strains are available upon request.

### Generation of the *ced-6* deletion allele

The *ced-6(hen38)* allele is a 768 bp deletion that removes the first and part of the second exon (flanking sequences are *atccgatccctcttccaact* and *atgcgtcgaaactccaaaag*). To create *ced-6(hen38)*, two cRNAs targeted to the *ced-6* locus were injected with the *dpy-10* cRNA and *dpy-10* repair oligonucleotide harboring the dominant *cn64* mutation. F1 rollers were cloned and screened for the deletion allele; F2 non-rollers were screened for homozygosity of the deletion allele.

### Transgenesis

Constructs used for cell specific expression of CED-1 and fluorescent proteins were generated using Gibson Assembly (NEB, E2611S), with cloning vector pPD95.75 used as the backbone. The 1 kb *col-10* promoter and 1.58 kb *klp-6* promoter were amplified from genomic DNA. *ced-1* cDNA was amplified from RNA isolated from N2 worms. *mCherry* was amplified from pHT101-mCherry (Addgene plasmid #61021). The *klp-6* promoter sequence and *ced-1* cDNA were assembled to create pTK8 (1.58 kb *klp-6* promoter::*ced-1*::*unc-54* 3’ UTR). The *col-10* promoter sequence and a PCR amplicon of the *ced-1* cDNA in pTK8 were assembled to create pTK9 (1 kb *col-10* promoter::*ced-1*::*unc-54* 3’ UTR). The *ced-1 cDNA* sequences in pTK9 and pTK8 were replaced with the *mCherry* sequences to create pTK16 (1 kb *col-10* promoter:: *mCherry*::*unc-54* 3’ UTR) and pTK18 (1.58 kb *klp-6* promoter::*mCherry*::*unc-54* 3’ UTR), respectively. pTK16 (25 ng/µl) and pTK18 (25 ng/µl) were independently injected into *ced-1(syb9348)* [CED-1::mNG] I; *him-5(e1490)* V young adult hermaphrodites along with genomic DNA (75 ng/µl) using the standard germline transformation protocol to generate extrachromosomal arrays. To create transgenic lines for cell specific rescue experiments, *ced-1(e1735)*; *mks-2(syb7299)*; *henSi21 him-5(e1490)* young adult hermaphrodites were injected. For CED-1 EVN rescue experiments, pTK8 (2 ng/µl), pTK18 (48 ng/µl) and genomic DNA (50 ng/µl) were injected; pTK18 (50 ng/µl) and genomic DNA (50 ng/µl) were injected together for the control. For CED-1 hypodermal rescue experiments, pTK9 (2 ng/µl), pTK16 (48 ng/µl) and genomic DNA (50 ng/µl) were injected; pTK16 (50 ng/µl) and genomic DNA (50 ng/µl) were injected together for the control. Four to six lines were isolated for each injection; we proceeded with the strains that had optimal fitness and the lowest expression level of the fluorescent proteins. For cell-specific rescue experiments, males with mCherry expression in the cells of interest (ex. either RnBs or hypodermis) were selected for analysis.

### EV imaging and analysis

Fourth larval stage (L4) males with *him-5(e1490)*, single copy transgenes expressing fluorescent protein-tagged EV cargos, and mutations of interest were staged with L4 hermaphrodites on the same plates 24 hours before EV imaging. Adult males were gently transferred onto an unseeded NGM plate to crawl free of the *E. coli* food source, then picked into 2.5 µl 20 mM levamisole on a 3% agarose pad on a glass slide. Animals were covered with high-performance cover glass (Zeiss, Item no. 474030-9020-000) that had been washed with ethanol in ultrasonic bath (40 kHz) for 30 minutes, rinsed three times in HPLC H_2_O for 10 minutes, and dried overnight to prevent contaminating signal from dust and other particles. EV images were collected with an Andor Dragonfly microscope and Zyla sCMOS detector at 63x using total internal reflection fluorescence (TIRF) microscopy; the angle of incidence was manually adjusted to achieve critical angle for each animal. Images of EVs released into the environment were taken 25-30 minutes after each animal was mounted. Mutant and relevant control strains were always imaged on the same day as there is day-to-day variation in EV shedding; images from at least three different days of imaging were pooled for analysis. EV images were quantified with the “Spot” function in Imaris software (Oxford Instruments), with the object size set to 0.350 μm in diameter.

### Cilia imaging and analysis

Age-synchronized adult males were immobilized in 2.5 µl 50 mM levamisole on 3% agarose pads on prewashed glass slides 24 hours post L4. Z-stack images were acquired with an Andor Dragonfly microscope (63x objective) in confocal mode and Zyla sCMOS camera. Mutant and relevant control strains were always imaged on the same day. Imaris (Oxford Instruments) was used to quantitate the volume, mean intensity and sum intensity of GFP-tagged proteins in the cilium proper and ciliary base as described.^19^ For experiments comparing mated males to virgin males, 20 late L4 males with or without 20 young adult hermaphrodites were staged on plates 24 h before imaging.

To analyze CED-1::mNG in the cilium, ciliary base, and distal dendrite of RnB neurons, the “multi plot” function in ImageJ (NIH) was used to plot fluorescence intensity along a 10 µm region of interest in dorsal RnBs (R1B and R5B) of splayed images. To exclude interference from CED-1::mNG in surrounding tissues, only 5 image slices (80 nm) selected based on the presence of the TZ in the focal plane were used for analysis. For each individual line scan, fluorescence intensity values were normalized to the maximum intensity in that line scan. The center of the transition zone, marked with MKS-2::mSc was used to align normalized fluorescence intensity measurements from different cilia.

### Mating behavior assessment

One day before analysis of mating behavior, L4 virgin males were picked onto new plates, separated from hermaphrodites. Mating plates made with low peptone (0.4g/L), which facilitates formation of a very thin bacteria lawn, were prepared with two droplets of 2.5 μl freshly cultured OP50. The next day, an individual adult male was placed on the mating plate with ten young adult hermaphrodites and behavior was recorded for 15 minutes with an AmScope (MU1000) camera and Amlite software. Male behaviors, including percent with contact response, ventral contact duration, turn numbers, and vulva stop duration were analyzed frame-by-frame in the recordings. Contact response was defined as a backward movement of the male along the hermaphrodite body after physical touch of the male tail on the hermaphrodite body. Upon reaching either the tail or the head of a hermaphrodite, the male tail must turn to continue scanning the hermaphrodite body for the vulva. Competed turns versus failed turns were defined based on whether the tail landed successfully on the other side of the hermaphrodite or not. Males that did not have physical contact with hermaphrodites (~10%) were not analyzed.

### Statistical analysis

GraphPad Prism version 10 was used for all statistical analyses and graphing. An Anderson-Darling normality test was used to evaluate all datasets. Following the normality distribution test, either a two-tailed Student’s *t* test or Mann-Whitney U test was used to compare two datasets; a one-way ANOVA was used when comparing four datasets. In all graphical analyses, *p< 0.05, **p< 0.01, ***p< 0.001, and ****p<0.0001. Details regarding the specific statistical test used for each experiment can be found in the figure legends.

## Notes

### Competing Interest Statement

The authors have declared no competing interest.

