## Supplemental Figures and Table for "*C. elegans* CED-1 acts in neurons to modulate ciliary protein abundance and extracellular vesicle shedding"

### Figure S1

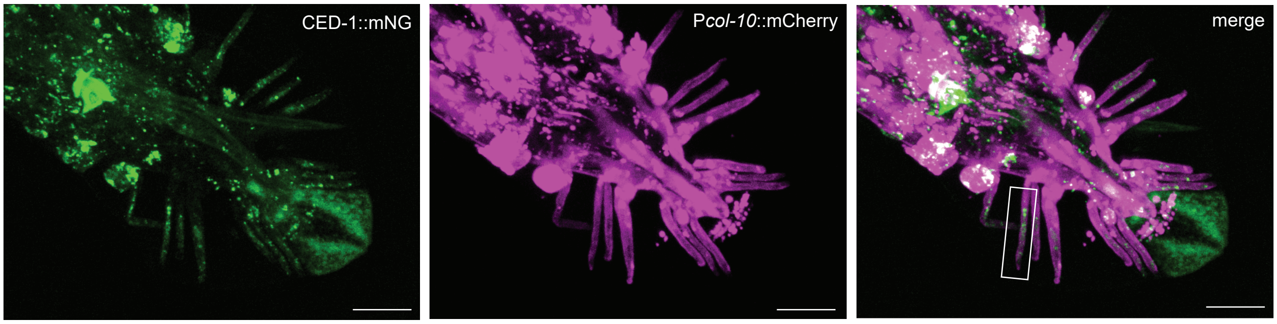

### Figure S2

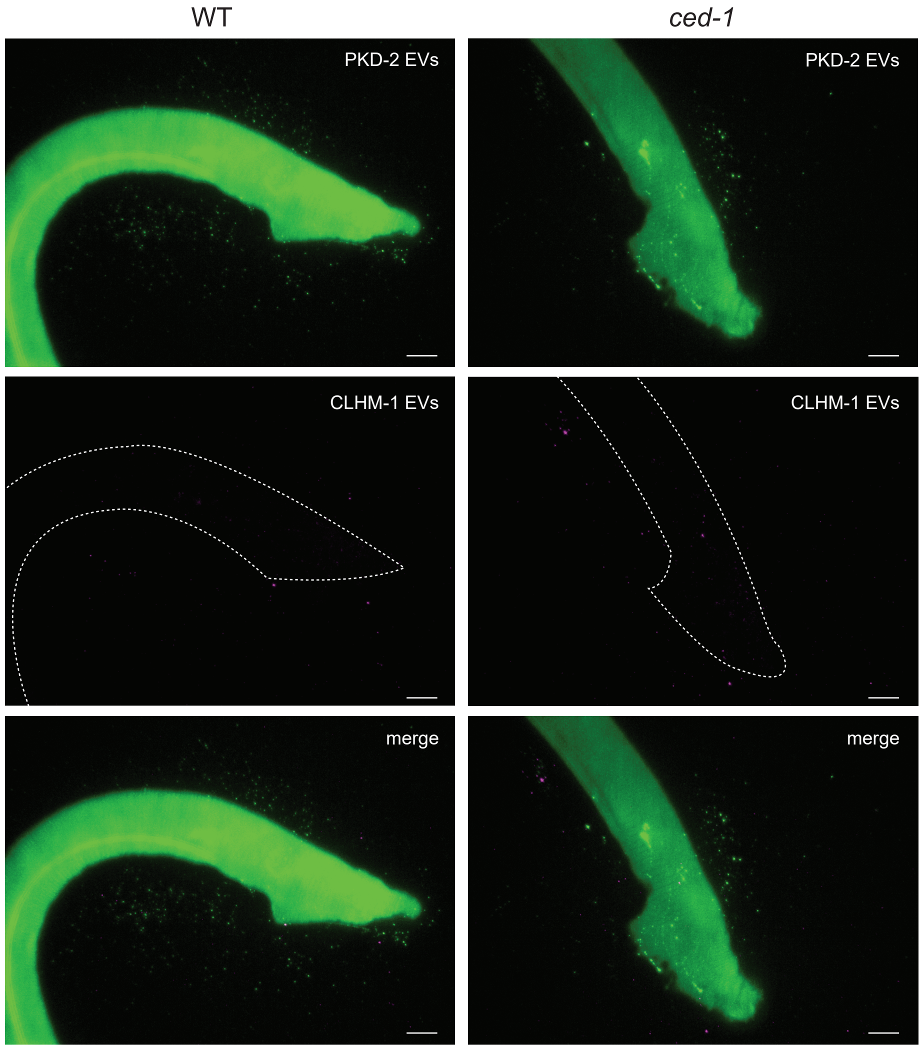

### Tabel S1

| Indexes of mating behaviors | WT (n=21) | *ced-1(e1735)* (n=23) | *ced-1(e1735)*;*Pklp-6*::mCherry (n=17) | *ced-1(e1735)*;*Pklp-6*::*ced-1* (n=22) |
|---|---|---|---|---|
| Contact response (%) | 79.11 (4.65) | 81.30 (3.89) | 75.01 (5.12) | 75.40 (5.11) |
| Ventral contact number | 4.61 (0.61) | 3.86 (0.66) | 2.71 (0.52) | 3.23(0.43) |
| Ventral contact duration (min) | 7.59 (0.77) | 6.47 (0.76) | 7.54 (0.84) | 6.83 (0.78) |
| Vulva stop number | 11.30 (1.58) | 10.86 (1.07) | 8.06 (1.26) | 7.46 (1.28) |
| Vulva stop duration (min) | 2.36 (0.36) | 3.20 (0.49) | 2.77 (0.54) | 2.51 (0.50) |
| Completed turns | 15.74 (2.44) | 12.81 (2.04) | 19.94 (5.38) | 14.91 (2.56) |
| Failed turns | 1.17 (0.29) | 2.76 (0.49)** | 3.35 (1.25) | 1.23 (0.42)[#] |
| Number of lost contact per min of ventral contact | 0.77 (0.12) | 0.80 (0.17) | 0.51 (0.13) | 0.66 (0.13) |
| Number of stops on vulva per min of ventral contact | 1.50 (0.17) | 1.98 (0.17) | 1.10 (0.12) | 1.11 (0.15) |
| Vulva stop duration per min of ventral contact (min) | 0.30 (0.04) | 0.46 (0.04)** | 0.35 (0.05) | 0.31 (0.05) |

The data are shown as mean +/- SEM; Mann-Whitney test; * $p<0.05$; ** $p<0.005$;
# $p=0.083$
